# Continuous embeddings of DNA sequencing reads, and application to metagenomics

**DOI:** 10.1101/335943

**Authors:** Romain Menegaux, Jean-Philippe Vert

## Abstract

We propose a new model for fast classification of DNA sequences output by next generation sequencing machines. The model, which we call *fastDNA*, embeds DNA sequences in a vector space by learning continuous low-dimensional representations of the *k*-mers it contains. We show on metagenomics benchmarks that it outperforms state-of-the-art methods in terms of accuracy and scalability.

## 1 Introduction

The cost of DNA sequencing has been divided by 100,000 in the last 10 years. With less than $1,000 to sequence a human-size genome, it is now so cheap that it has quickly become a routine technique to characterize the genome of biological samples with numerous applications in health, food or energy. Besides the genome, many techniques have been developed to measure other molecular informations using DNA sequencing, e.g., gene expression using RNA-seq, protein-DNA interactions using ChiP-seq, or 3D structural informations using Hi-C, to name just a few. In short, DNA sequencing is the swiss army knife of modern genomics and epigenomics. As a consequence, the rate of production of DNA sequences has exploded in recent years, and the storage, processing and analysis of these sequences is increasingly a bottleneck.

While new, so-called *long-read* technologies are under active development and may become dominant in the future, the current market of DNA sequencing technologies is dominated by so-called *next-generation sequencing* (NGS) technologies which break long strands of DNA into short fragments of typically 50 to 400 bases each, and “read” the sequence of bases that compose each fragment. The output of a typical DNA sequencing experiment is therefore a set of millions or billions of short sequences, called *reads*, of lengths 50 ~ 400 in the {*A, C, G, T*} alphabet; these billions of reads are then automatically processed and analyzed by computers to get some biological information such as the presence of particular bacterial species in a sample, or of a specific mutation in a cancer.

Standard pipelines to process the raw reads depend on the target applications, but typically involve discrete operations such as aligning them to some reference genome using string algorithms. In this paper, we investigate the feasibility of directly representing DNA reads as continuous vectors instead, and replacing some discrete operations by continuous calculus in this embedding.

To illustrate this idea, we focus on an important application in metagenomics, where one sequences the DNA present in an environmental sample to characterize the microbes it contains [21, 4]. An important problem in metagenomics is *taxonomic binning*, where each of the billions of sequenced reads must be assigned to a species, given a database of genomes characteristic of each species considered [16]. Standard computational approaches for taxonomic binning try to align each read to a reference sequence database with sequence alignment tools like BLAST [5] or short read mapping tools such as BWA [14] or BOWTIE [10]. However, the computational cost of these techniques becomes prohibitive with current large sequence datasets. Alternatively, compositional approaches rephrase the problem as a multiclass classification problem, and employ machine learning methods such as a naive Bayes (NB) classifier [23, 19] or a support vector machine (SVM) [17, 20, 22] after representing each read by the vector of *k*-mer^1^ counts it contains. Interestingly, [22] showed that compositional approaches can be competitive in accuracy with alignment-based methods, while maintaining a computational advantage, by using large-scale machine learning approaches. However, a good accuracy is only achieved with *k*-mers of length at least *k* = 12, corresponding to representing each read as a sparse vector in *N* = 4*^k^* dimensions. This representation raises computational challenges both at training time ([22] push VowPal Wabbit to its limit) and at test time (a *N* × *T* matrix of weights must be stored to model *T* species).

In this work, we propose to extend state-of-the-art compositional approaches by embedding the set of DNA reads to ℝ^*d*^, with *d* ≪ *N*. For that purpose, we still extract the *k*-mer composition of each read, but replace the *N*-dimensional one-hot encoding of each *k*-mer by a *d*-dimensional encoding, optimized to solve the task. This approach is similar to, e.g., the `fastText` model for natural language sequences of [7, 3] or `word2vec` [18], with a different notion of *words* to embed, and a direct optimization of the classification error to learn the representation. This can reduce the memory requirements to store the model and accelerate classification time when *d* < *T*, since the *N* × *T* matrix of weights is replaced by a *N* × *d* matrix of embeddings, and a *d* × *T* matrix of weights.

After presenting in more detail the model and its optimization, we experimentally study the speed/performance trade-off on metagenomics experiments by varying the embedding dimension, and demonstrate the potential of the approach which outperforms state-of-the-art compositional approaches.

## 2 Method

### 2.1 Embedding of DNA reads

Given the alphabet of nucleotides *𝒜* = {*A, C, G, T*}, a *DNA read* of length *L* ∈ ℕ* is a sequence x = *x*_1_ *… x_L_* ∈ *𝒜^L^*. Depending on the sequencing technology, ***L*** is typically in the range 50 *∼* 400, and we fix *L* = 200 in the experiments below. For any 1 ≤ *a* ≤ *b* ≤ *L* we denote by x_[*a,b*]_ = *x_a_x_a_*_+1_ *… x_b_* the substring of x from position *a* to *b*. For any *d* ∈ ℕ*, an *embedding* of DNA reads to ℝ*^d^* is a mapping Φ: *𝒜^L^ →* ℝ*^d^* to represent each read x ∈ *𝒜^L^* by a vector Φ(x) ∈ ℝ^*d*^, which can then be used for subsequent classification tasks.

For a given *k* ∈ ℕ, (the *k*-spectral embedding represents a sequence by its *k*-mer profile [11]: it is an embedding 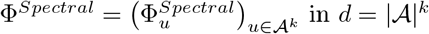 dimensions indexed by all strings of length *k*, where for any such string *u* ∈ *𝒜^k^* one defines:

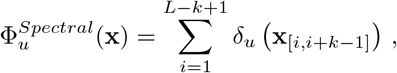

where *δ_u_*(*v*) = 1 if *u* = *v*, 0 otherwise. The *k*-spectral encoding of DNA reads is used in state-of-the-art compositional approaches to assign reads to species with machine learning techniques [23, 19, 17, 20, 22].

Given *d* ∈ ℕ^*^ and a *N*×*d* matrix 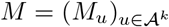 associating a vector *M_u_* ∈ ℝ*^d^* to each *k*-mer *u*∈*𝒜^k^*, we now consider a *d*-dimensional embedding Φ*^M^* of DNA reads by summing the vectors associated to the read’s *k*-mers:

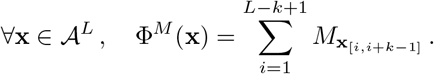

In matrix form, one easily sees that the *d*-dimensional embedding Φ*^M^* can be obtained from the *k*-spectral representation by the formula:

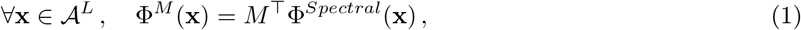

showing in particular that the *k*-spectral embedding is a particular case of Φ*^M^* by taking *d* = *N* and *M* = *Id*. Changing *M* allows to create correlations between *k*-mers in the embedding space. For example, the (*k, m*)-mismatch kernel [12] also corresponds to an embedding Φ*^M^* with *d* = *N*, but where *M_u,v_* = 1 when the Hamming distance between *u* and *v* is at most *m*, 0 otherwise. Changing *d* further allows to vary the dimension of the embedding, which can not only be beneficial for memory and computational reasons, but also help statistical inference by reducing the number of parameters of the embedding.

### 2.2 Learning the embedding

While several existing embeddings such as the *k*-spectral or (*k, m*)-mismatch embeddings correspond to Φ*^M^* for specific matrices *M*, we propose to “learn” *M* as part of the overall classification or regression task that must be solved. In our metagenomics problem, this is a multiclass classification problem where each of the *T* bacterial species is a class and each read must be assigned to a class. Given an embedding Φ*^M^*, we consider a linear model of the form:

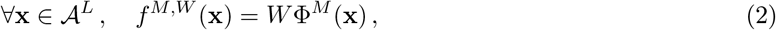

where *W* ∈ ℝ^*T* × *d*^ is a matrix of weights, and the prediction rule:

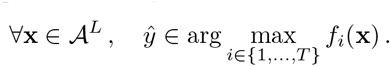

To learn the embedding *M* and the linear model *W*, we assume given a training set of examples 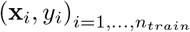 where *x_i_* ∈ *𝒜^L^* and *y_i_* ∈ {1, …, *T*}, and numerically minimize an empirical risk:

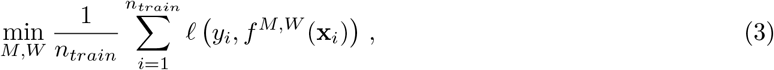

where for the loss *ℓ* we choose the standard cross-entropy loss after transforming the scores to probabilities with the softmax function:

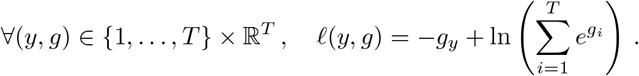

We solve (3) by stochastic gradient descent (SGD). Note that when *d* < *T*, the problem is usually non-convex and SGD may only converge to a local optimum.

Combining (1) and (2), we further notice that for any embedding *M* and weights *W*,

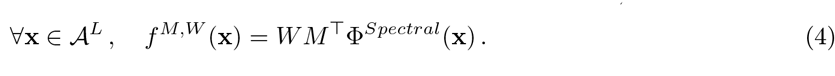

This clarifies that *f ^M,W^* boils down to a linear model in the *k*-spectral representation, with a weight matrix *WM*^Τ^ of rank at most *d*. When *d* < min(*N, T*) this creates a low-rank regularization that can be beneficial for statistical inference, in addition to reducing the memory footprint of the model and speeding up the prediction time.

### 2.3 Implementation

We implemented the model (3) by modifying the `fastText` open-source library [7, 8], which involves a similar model with *k*-mer embedding for natural language. There are some important differences between DNA reads and standard NLP applications, though: (i) One is that the concept of a “word”, space-delimited groups of letters, does not exist in DNA sequence data. Hence we resort to a distributed representation of overlapping *k*-mers only, and not to words as in `word2vec` or `fastText`. (ii) Second is the number of training examples to be seen by the model is very large: up to 5 × 10^9^ if we were to achieve full coverage of the *large* database below, for example. (iii) Third, the vocabulary is different, as it is of known size (4^*k*^) and is densely represented for relatively small values of *k*. For greater values of *k* than those considered in this paper, *k*-mers become rare and individual long *k*-mers can become discriminative. This is in fact used by some other compositional algorithms such as Kraken [24].

For these reasons we rewrote part of the `fastText` software to extract overlapping *k*-mers rather than words. As appropriate for (iii), the *k*-mer embeddings are stored in a fixed-size table of dimension (4*^k^, d*), each row corresponding to the vector of a different *k*-mer. For a given *k*-mer **b** = *b*_1_ *… b_k_* ∈ *𝒜^k^*, the index of its corresponding row in *M* is 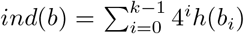, where *h*(*A*) = 0, *h*(*C*) = 1, *h*(*G*) = 2 and *h*(*T*) = 3. This allows to quickly find *ind*(**x**_*i*+1:*i*+*k*_) from *ind*(*i*: *i* + *k−*1) as explained for example in [22]. To address (ii) we generate random fragments on the fly directly from the full genomes, rather than reading text samples line by line as in `fastText`.

### 2.4 Regularization with noise

As a form of regularization, we also add random mutations in the training fragments. When reading the fragment from the reference genome, nucleotide by nucleotide, we introduce a chance *r* of replacing the nucleotide by a random one (equally distributed on {*A, C, G, T*}). This is akin to the dataset augmentations commonly used in image classification tasks, and in our case promotes the nearby embeddings of *k*-mers similar in terms of Hamming distance.

## 3 Experiments

### 3.1 Data

We test our model on two benchmarks proposed by [22]: a small one, useful mostly for parameter tuning, and a large one. Both benchmarks involve a *training* database of genomes organized by species, and a *validation* set of genomes coming from the same species as the training database, but from different strains. Reads are randomly sampled, with or without noise, from the validation set of genomes, and the goal is to predict, for each read, from which species it comes from. The `small` database contains 356 complete genomes, belonging to 51 species of bacteria; its validation set is composed of 52 genomes, belonging to the same 51 species. The `large` database contains 2,961 genomes belonging to 774 species, which is closer to real-life situations. The validation set is composed of 193 genomes, each from a separate species.

For both training and testing, reads of length *L* = 200 are extracted from the genomes. The validation datasets are built by extracting fragments such that their coverage – the average number of times each nucleotide is present – is 1. This amounts to a total of 134, 319 validation samples for the *small* database, and ~ 3.5M samples for the *large* database. The machine learning-based models are trained on reads sampled from the reference genomes and their known taxonomic labels, while alignment-based methods simply align validation reads to the training reference genomes. To account for the fact that DNA is double-stranded and that when a read is sequenced it can come from any of the two strands, which are reverse-complement to each other, we systematically add the reverse-complement of each read with the same label at training time.

In addition, we consider several *noisy* validation sets as in [22], where each fragment sampled from a genome is modified to mimick sequencing errors of actual sequencing machines, in particular substitutions, insertions and deletions of nucleotides. We use the specially-designed `grinder` software [1] to simulate 3 new sets of validation reads. The *Balzer* validation set is simulated with a homopolymeric error model, designed to emulate the Roche 454 technology [2]. The *mutation-2* and *mutation-5* sets are simulated with the 4th degree polynomial proposed by [9] to study general mutations (insertion/deletions and substitutions). The median mutation rates for these simulated reads are 2% and 5%, respectively. *Balzer* and *mutation-2* are meant to contain a realistic proportion of errors, and *mutation-5* is added as a more challenging set.

### 3.2 Reference methods

We compare our method, which we call `fastDNA` in the rest of the text, to two other strategies. One is the BWA-MEM sequence aligner [13] and the other is the linear SVM classifier on the *k*-spectral representation, implemented using the Vowpal Wabbit software in [22]. We name the latter method VW in the rest of the paper. We follow exactly the same configurations as [22] for both methods.

### 3.3 Small dataset

#### 3.3.1 Memory footprint

As the embeddings matrix *M* is loaded in memory both for training and classification, `fastDNA` models have a significant memory footprint. [7] discuss various strategies to reduce it. The vocabulary cannot be pruned for our range of *k* as the *k*-mers are densely distributed, so the dimensions of *M* are fixed. The size of *M* in memory can however be reduced by quantization. We use Product Quantization (PQ, [6]), with the option to cluster separately the vector norms and directions (option `qnorm`). The linear layer is then retrained to account for the change in the embeddings. This compresses the model size by almost an order of magnitude without noticeably impacting the performance.

While PQ can make deploying and classifying more accessible, training the model still requires the full embedding matrix, as its quantized version is not trainable. We restrain our parameter choices (*k* and *d*) to models that fit on 64GB machines. The dimension of the embedding table (4^*k*^) is encoded on 32-bits, which further limits the value of *k* to *k*_max_ = 15. The largest models we consider are *k* = 13, *d* = 100 and *k* = 15, *d* = 10. Their memory footprints are available in figure 1.

**Figure 1:**
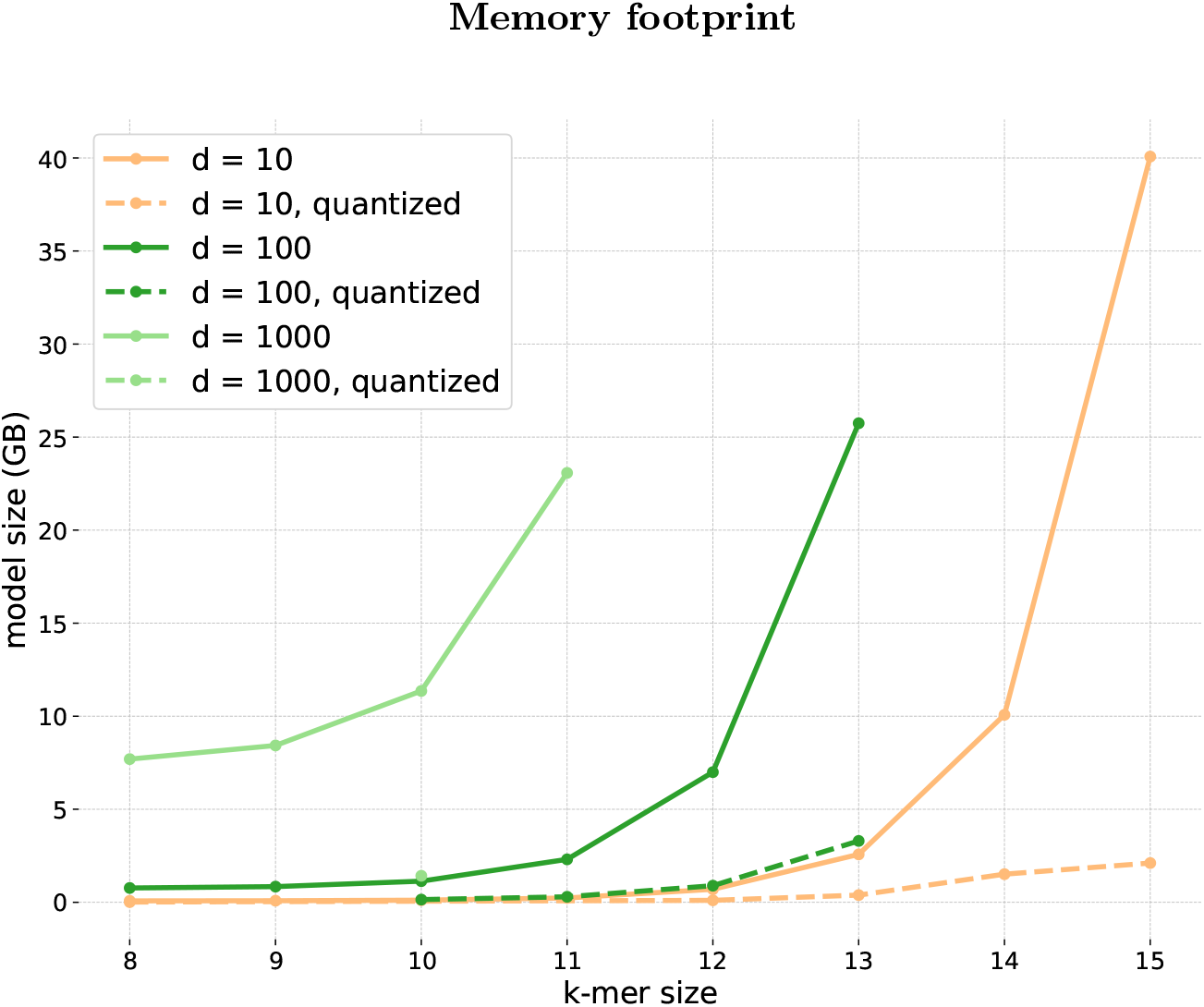
Memory requirement of `fastDNA` models as a function of *k*-mer sizes. The embedding dimensions *d* shown are 10, 100 and 1000. The reported value is the size in GB of the model binary, the minimal size required both in RAM to train and load the model, and on disk to save it.

Contrary to VW, model size is completely determined by the user and does not depend on the number of possible classes *T* or on the vocabulary of the training database, which can become an advantage when *T* is large.

#### 3.3.2 Coverage

The model is trained by picking a position at random in the reference genomes and using the 200-bp read starting from that position as a sample. One epoch of training consists of drawing enough random reads to cover each nucleotide of the reference genomes once on average (coverage of 1). We found that models with no training noise had converged by 50 epochs, and those with training noise benefited from extra epochs.

The results in this paper were obtained by training with a fixed number of epochs 50, and a learning rate of 0.1, chosen by a standard grid-search.

#### 3.3.3 Performance

One first result from figure 2, is that increasing the embedding dimension *d* above 50 brings no added benefit in classification quality, at the cost of larger prediction times and memory footprint. This could in theory be expected from (4). Once *d* is greater than the number of classes, the matrix *WM*^Τ^ has maximal rank *T*, so the model is virtually the same as the standard “Bag of Words” model VW. The differences observed between the two are likely due to different optimization procedures and implementations.

**Figure 2:**
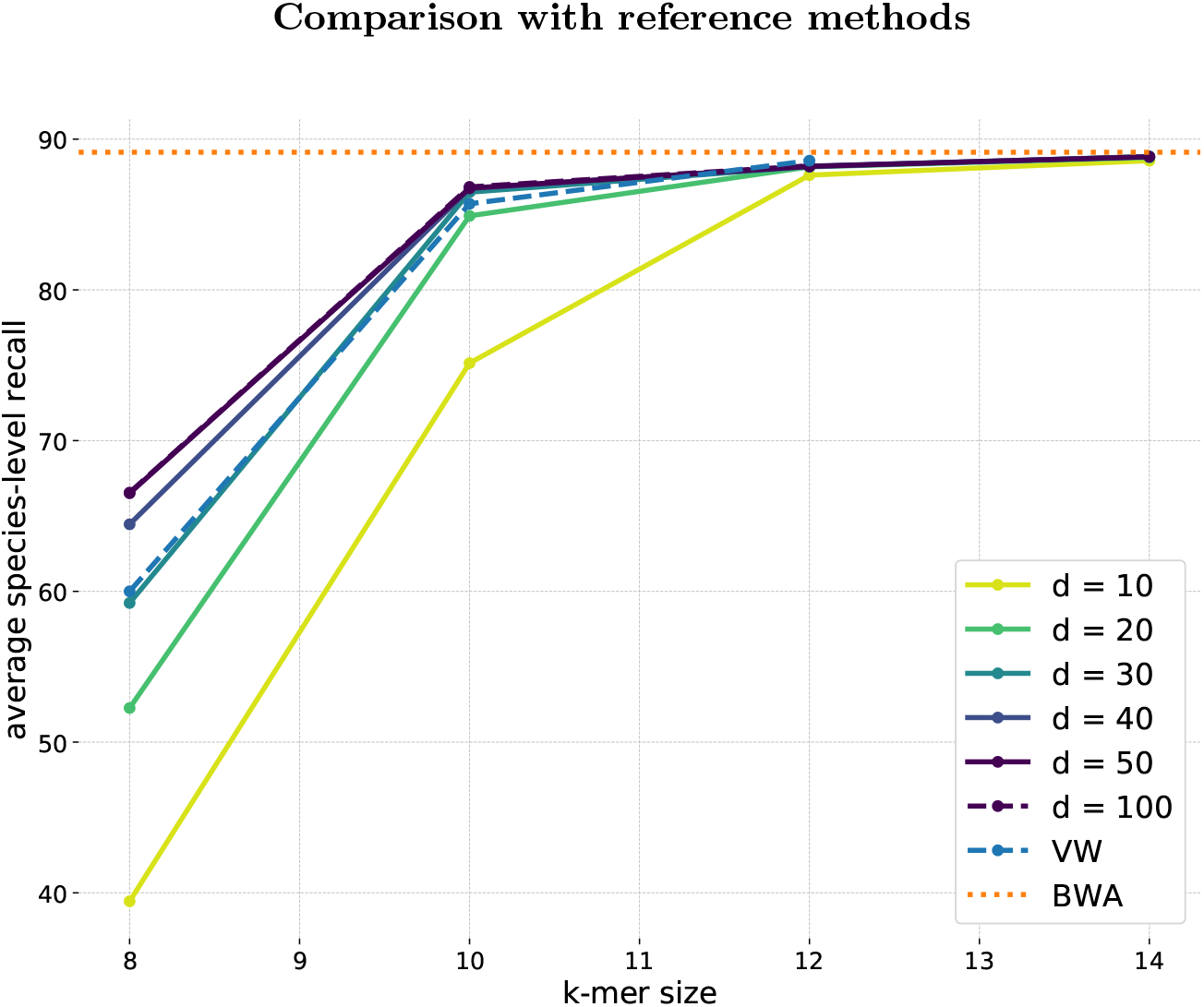
Comparison between `fastDNA` and reference methods on the *small* dataset. This figure shows the average species-level recall obtained by `fastDNA` trained for 50 epochs and a learning rate 0.1, for different values of *k* and *d*. The results are compared with VW for different values of *k* and an alignment-based approach BWA-MEM

Excessively lowering the dimension *d* under *T* harms the performance, especially for shorter *k*-mers. However, the gap between models *d* = 10 and models *d* = 50 vanishes as *k* increases, suggesting that – provided a sufficient vocabulary size – projecting *k*-mers to a lower-dimensional space comes at little cost. Furthermore, models with longer *k*-mers but smaller dimension can achieve the same performance as a model with shorter *k* and greater dimension. The model *k* = 14, *d* = 10 has the same performance as *k* = 12, *d* = 50.

Finally, classification performance for values of *k* greater than 12 is competitive with alignment-based method BWA, which confirms machine learning approaches can be relevant for this problem of taxonomic binning.

### 3.4 Large dataset

We report the classification performance, measured by average species-level recall and precision of `fastDNA` on the validation sets described in 3.1. The influence of the training noise is shown in figure 3.

**Figure 3:**
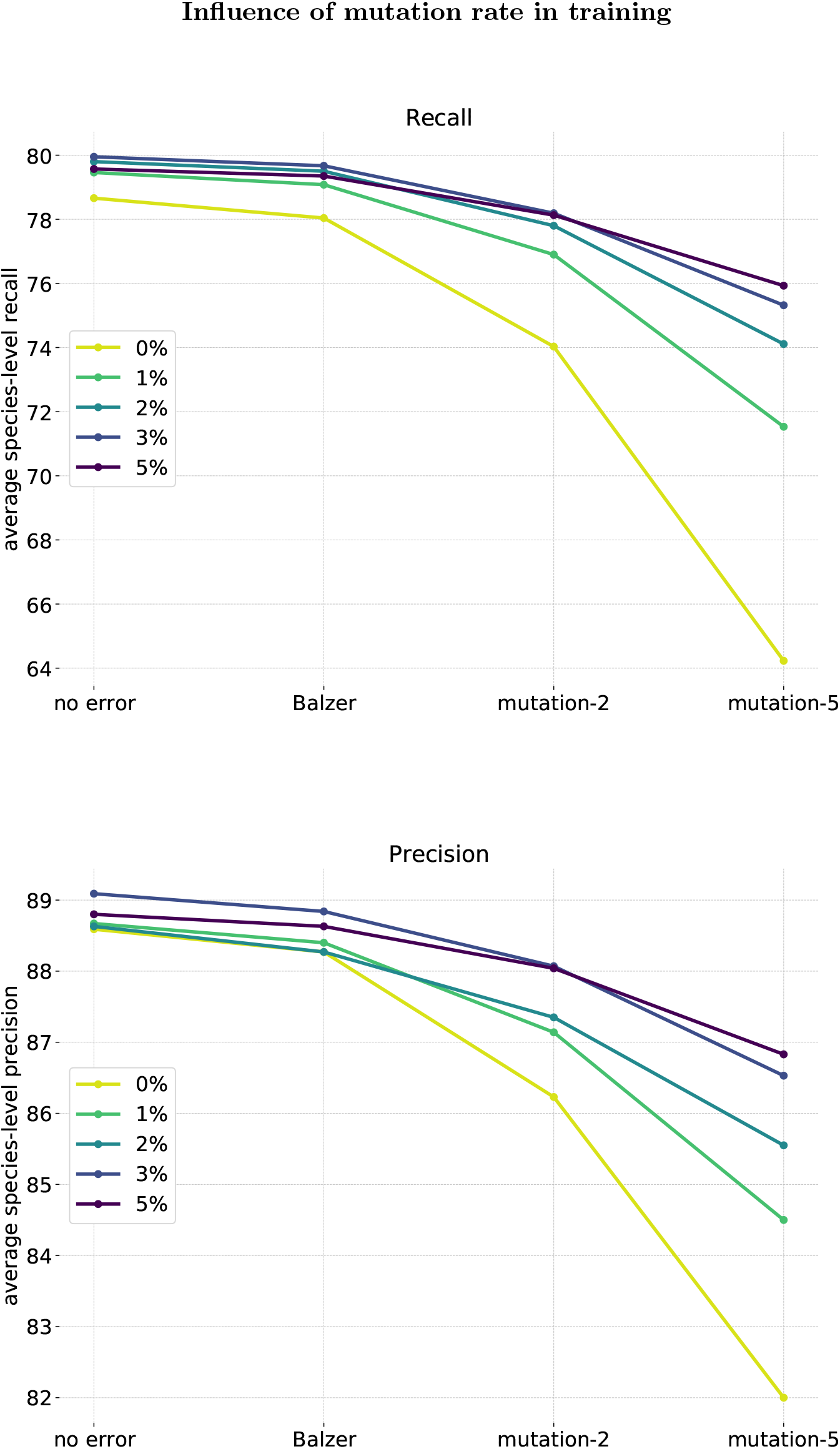
Performance on the large dataset of `fastDNA` trained with different mutation rates. *k*-mer size and embedding dimension *d* are 13 and 100, respectively. The classification quality is measured on test sets generated with different sequencing error models.

**Figure 4:**
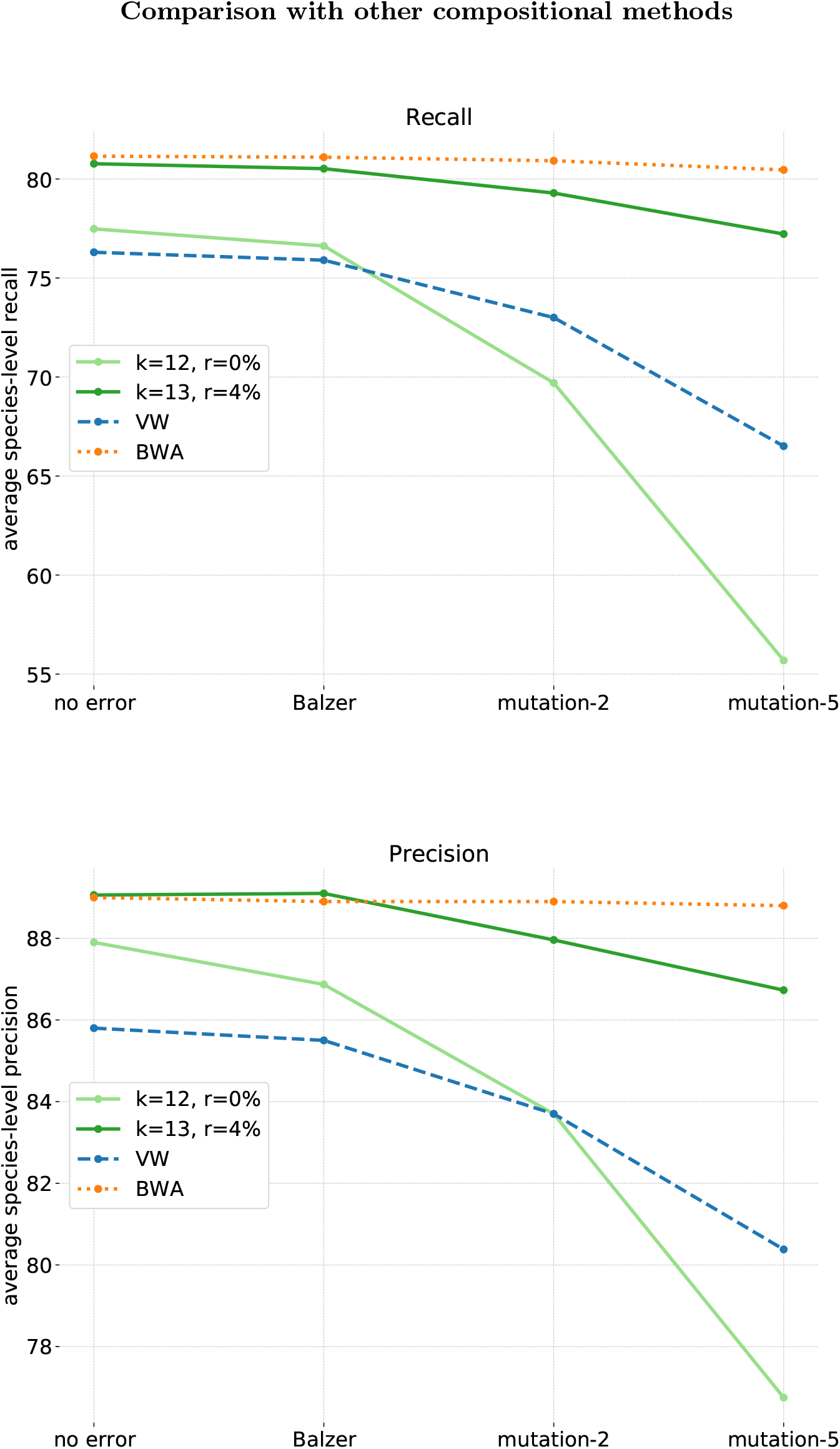
Comparison between `fastDNA` and reference methods on the *large* dataset. This figure shows the average species-level recall and precision obtained by fastDNA, VW and BWA on the different validation sets. We show both the best configuration for `fastDNA` (*k* = 13, *d* = 100, *r* = 4%) and a similar one to VW (*k* = 12, *d* = 100, *r* = 0%) for a fair comparison.

As could be expected, greater levels of sequencing noise in the validation sets lead to degraded performances. Adding random mutations to the training reads curbs this effect. The greater the mutation rate is, the more robust the model becomes to higher levels of sequencing noise. Somewhat more surprisingly, a certain range of mutation rates also increases the performance on the validation reads with no sequencing noise. The regularization induced by these artificial errors is therefore beneficial for both sequencing noise and intra-species heterogeneity. We found that the best rates of mutation were between 2 and 5%. The models trained with these levels of noise are better all-around than their no-noise counterparts.

We compare the performance of `fastDNA` against that of VW and BWA in 4. For small levels of sequencing errors, `fastDNA` is competitive with BWA. Greater levels in sequencing noise widens the gap between the two, as BWA is very robust to sequencing noise, dropping less than 1% for *mutation-5*.

### 3.5 Classification speed

Speed is of critical importance in taxonomic binning and is the main motivation behind exploring machine learning techniques. Classifying a read with `fastDNA` can be separated in two parts. It first reads the sequence, computes the indices of the *k*-mers contained in the read and computes the read embedding by summing the *k*-mer embeddings. This step is of complexity 𝒪 (*dL*), where *L* is the read length (constant in our experiments). Second, class probabilities are computed by applying the linear classifier, this step is of complexity 𝒪 (*dT*). `fastText` and `fastDNA` offer a different loss function, the hierarchical softmax, that reduces this step to 𝒪 (*d* log(*T*)), which can become useful in the case of very large *T*. To this per-read time-complexity must be added a fixed overhead, the time necessary for the model to be loaded from disk, of complexity 𝒪 (*dN* + *T*), where *N* = 4^*k*^ is the vocabulary size. Due to the large values of 4^*k*^, this can make up a significant portion of the total time. Moreover, we observe in practice a longer memory access time for larger vectors. These are the two reasons for the time gap observed between classification with models of same embedding dimension *d* but different *k*.

The total time complexity for predicting a dataset with *n* samples is therefore 𝒪(*n*(*dL* + *dT*) + *dN*).

With reads of constant length, `fastDNA` and VW classify reads indiscriminately of their content, and therefore yield equal classification times across the validation sets. On the other hand, BWA’s speed degrades with the sequencing noise, which is easily explained. BWA searches iteratively on the number of mismatches *z* and stops once it gets a hit. It will therefore be slower if there are more mismatches between the testing and reference data.

We show in figure 5 the classification speeds measured for the *large* dataset. The values reported are for a single CPU (Intel Xeon E5-2450 v2 −2.5GHz). We show `fastDNA` for *d* = 100 and *k* = 12, 13 and VW. `fastDNA` and VW have similar classification times of ~6, 7 10^3^ reads per minute. As remarked in [22], compositional approaches offer systematically better prediction times than BWA, with improvements of 2−9 ×. This speed improvement increases with the mismatch between predicted sequences and reference genomes, therefore with both sequencing noise and intra-species variations.

**Figure 5:**
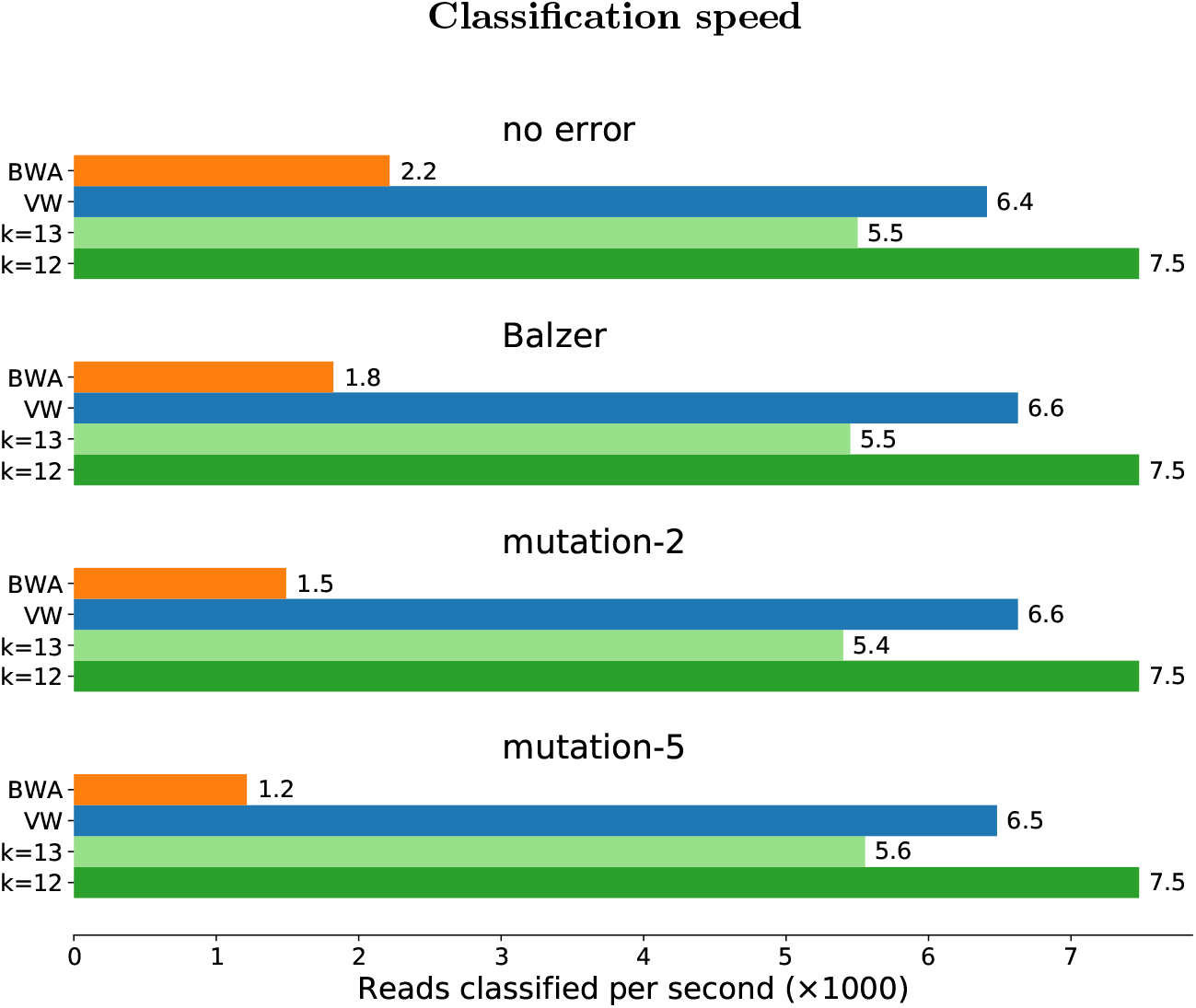
Comparison between `fastDNA` and reference methods on the *large* dataset. This figure shows the average classification speed of the methods on the different test sets. Two versions of `fastDNA` are shown, one with *k*-mer size 12, the other with *k*-mer size 13. Both have embedding dimension *d* = 100. The four test sets used were simulated with different sequencing error models.

## 4 Conclusion

We demonstrated that learning a low-dimensional representation of DNA reads based on their *k*-mer composition is feasible, and outperforms state-of-the-art compositional approaches that work directly on the high-dimensional, *k*-spectral representation of DNA sequences. Controlling the dimension *d* of the embedding allows to consider longer *k*-mers for a given memory footprint. As other compositional methods, `fastDNA` is significantly faster than alignment-based methods, and is well adapted to classification into many classes.

There are two immediate possible extensions of this work. One is to use a more realistic error model than uniform substitutions for training, to better mimick the expected noise in the data. The other is to extend the notion of “word” from contiguous *k*-mers to gap-seeded *k*-mers or Bloom filters [15], hopefully capturing longer-range dependencies. In terms of applications, assigning RNA-seq reads to genes to quantify their expression can also be formulated as a classification problem with typically *T* ∼ *k* classes, and may be well adapted to `fastDNA` as well.

The source code is freely available and published on github https://github.com/rmenegaux/fastDNA, along with scripts to reproduce the presented results.

## Footnotes

1 A *k*-mer is a contiguous subsequence of *k* letters.

